# A confound-diagnostic toolkit for in silico perturbation with single-cell foundation models

**DOI:** 10.64898/2026.08.04.732812

**Authors:** Ru Qiu, Max Mingqian Zhao

**Affiliations:** State Key Laboratory of Ophthalmology, Zhongshan Ophthalmic Center, Sun Yat-sen University, Guangdong Provincial Key Laboratory of Ophthalmology and Visual Science, Guangzhou 510060, Guangdong, China; Department of Electrical and Computer Engineering, The University of Texas at San Antonio, San Antonio, TX 78249, USA; Health Careers High School, Northside Independent School District, San Antonio, TX 78229, USA

**Keywords:** single-cell foundation models, in silico perturbation, Geneformer, Perturb-seq, confound diagnostics, tokenization coverage

## Abstract

Deleting a gene token from a cell’s input sequence offers a convenient native strategy for in silico perturbation, but the resulting embedding delta may not represent a biological knockout response. Apparent effects can instead reflect gene identity, universal responsiveness, limited tokenization coverage, library-size contamination, or circular state scoring. Here, we present a confound-diagnostic framework combining held-out increment testing, responsiveness adjustment, coverage gating, library-size diagnostics, and de-circularized state-shift analysis, together with a numerically matched reimplementation of frozen Geneformer’s perturbation engine. Across Frangieh and Replogle datasets and linear and nonlinear readouts, the native embedding delta provided no reproducible held-out improvement beyond gene identity. Signal-injection calibration showed that the test detected injected residual signal, whereas native increments remained below its detection floor. Matched controls traced apparent positives to raw-count library-size structure, broad responsiveness, and self-referential scoring, while coverage constrained perturbation applicability and estimate stability without establishing biological specificity. This model-adaptable framework helps determine when foundation-model perturbation readouts warrant biological interpretation.

**Motivation:** Foundation-model in silico perturbation could predict perturbation effects when matched experimental data are unavailable. However, in zero-shot settings, embedding-derived responses may reflect gene identity, universal responsiveness, tokenization limits, library-size artifacts, or circular state scoring rather than biological knockout effects. We therefore developed a reusable confound-diagnostic framework that applies matched controls to test whether native perturbation readouts contain information beyond these confounds and warrant biological interpretation.

## Introduction

Single-cell foundation models, including Geneformer^1^, scGPT^2^ and scFoundation^3^, have created new opportunities for learning transferable representations of genes and cells from large-scale transcriptomic data. One proposed application is zero-shot in silico perturbation. In tokenized models such as Geneformer, this can be implemented by removing a target-gene token from a cell’s input sequence and interpreting the resulting change in model embedding, referred to here as the embedding delta, as a predicted knockout response. This native token-deletion strategy is attractive because it could, in principle, estimate perturbation effects when matched experimental perturbation data are unavailable. It also differs from network-inference and supervised perturbation-response methods, such as CellOracle^4^, scGen^5^, and GEARS^6^, which typically require additional regulatory information, chromatin data, or perturbation training datasets.

Despite this promise, the biological validity of native foundation-model perturbation readouts remains unresolved. Recent benchmarking studies have reported limited zero-shot utility on general single-cell tasks^7^ and on perturbation-specific evaluations^8,9^, modest or inconsistent gains over simple baselines^10,11^, and substantial sensitivity to metric choice, normalization strategy, baseline definition, and target readout^12^. These findings indicate that apparent predictive performance alone is insufficient to establish perturbation-specific biological signal. Rigorous evaluation therefore requires determining whether an embedding-derived perturbation readout provides information beyond confounding structure in the model representation, input data, and evaluation design.

Several confounding factors can inflate apparent perturbation readouts, making them appear informative without necessarily reflecting perturbation-specific biological signal^13^. Gene identity and baseline expression structure can make affected genes predictable even in the absence of a perturbation-induced embedding delta. Highly expressed or broadly connected genes can appear responsive across many perturbations, producing non-specific saliency patterns. Differential-expression targets derived from raw counts can be distorted by library-size variation. In tokenized models, a nominal knockout may have limited meaning when the target gene is absent from most truncated input sequences. Cell-state-shift analyses can also become circular when a deletion is evaluated against a state axis partly defined by that same deletion. These failure modes are mechanistically distinct and require matched diagnostic controls.

The central methodological need is therefore a framework that evaluates whether native foundation-model perturbation readouts remain informative after explicit control for known confounds. Such a framework should test incremental value beyond gene identity, account for universal responsiveness in affected-gene saliency, verify that the perturbation is applied in a sufficient fraction of cells, protect differential-expression targets from library-size artifacts, and avoid self-referential scoring in cell-state projections. A confound-aware workflow would provide practical safeguards for interpreting current token-deletion readouts and for evaluating future foundation-model perturbation methods.

Here, we present a reusable confound-diagnostic workflow for evaluating native foundation-model in silico perturbation, using frozen Geneformer token deletion as a case study. The workflow integrates held-out increment testing, universal-responsiveness adjustment, tokenization-coverage gating, library-size diagnostics, and de-circularized state-shift analysis. A numerically matched reimplementation of Geneformer’s native perturbation response enables controlled coverage-to-estimate analyses. Across the Frangieh and Replogle datasets and across linear and nonlinear readout models, native embedding deltas showed no reproducible improvement beyond gene identity. Signal-injection calibration confirmed that the increment test was sensitive to injected residual signal, and matched controls attributed apparent positive findings to library-size structure, universal responsiveness, and self-referential scoring. Tokenization coverage constrained perturbation applicability and estimate stability but did not establish biological specificity. Although the empirical analysis is limited to frozen Geneformer, the framework is broadly applicable for determining when foundation-model perturbation readouts warrant biological interpretation.

## Results

### A numerically matched reimplementation enables confound-controlled testing of embedding-delta signal

We first established a reproducible reimplementation of Geneformer’s native in silico knockout response. In this token-deletion procedure, the target-gene token is removed from each cell’s input sequence, and the resulting change in gene or cell embeddings is summarized as the native perturbation response. We reimplemented the per-cell gene cosine-similarity response to target-gene deletion, referred to as path-D, and validated it against the official Geneformer InSilicoPerturber output. Path-D reproduced the official output to numerical precision, with Pearson’s r>0.999999 and a maximum absolute error of 3×10^−7^.

This matched reimplementation provided the computational basis for the confound-diagnostic workflow. Cached per-cell path-D responses enabled controlled resampling while holding the model checkpoint, tokenization, target gene, and perturbation operator fixed. The workflow incorporated held-out increment testing against a gene-identity baseline, universal-responsiveness adjustment, tokenization-coverage gating, library-size diagnostics, and de-circularized cell-state-shift analysis (**Figure 1**; **Table 1**).

**Figure 1.**
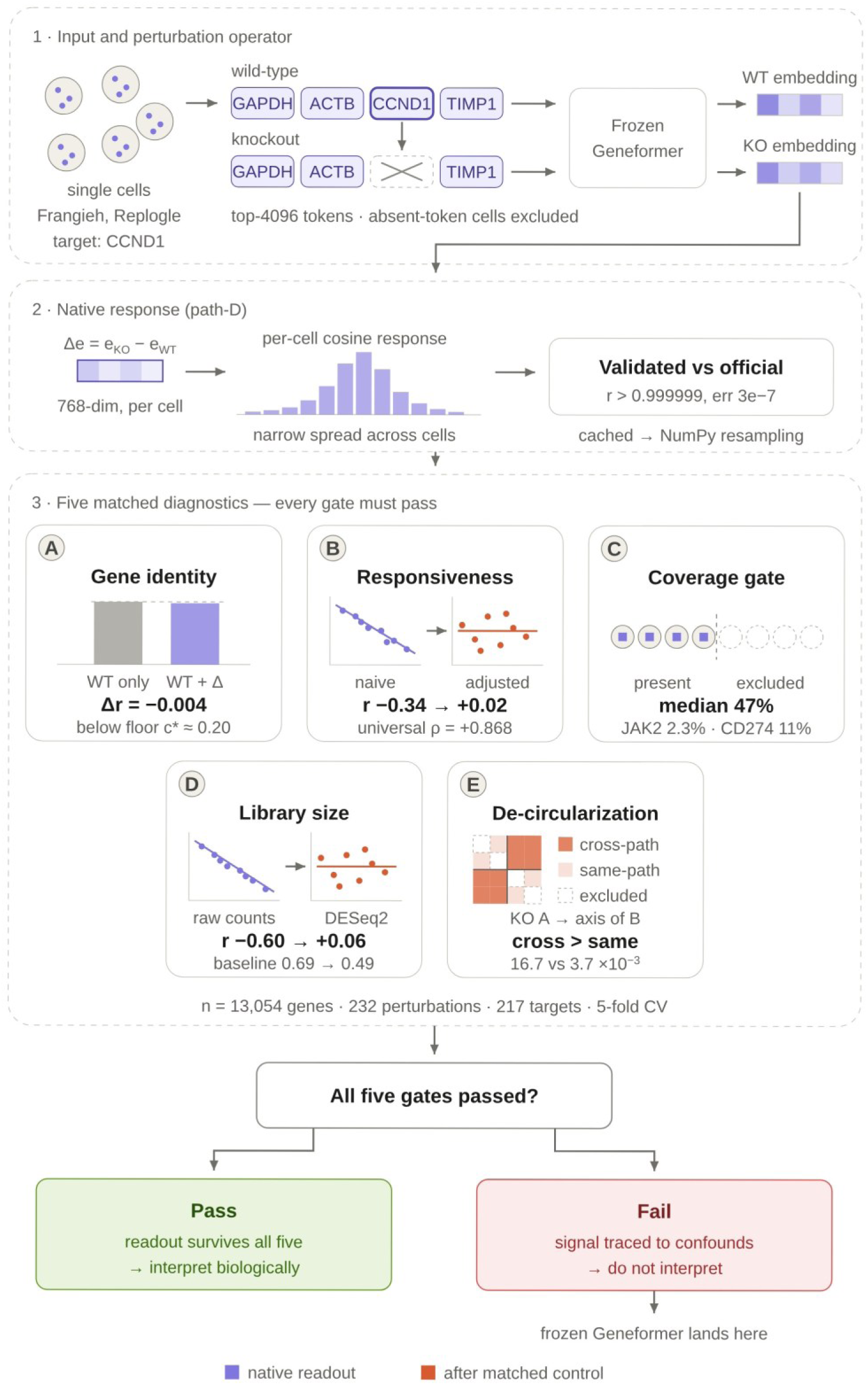
Overview of the confound-diagnostic workflow. Native Geneformer token-deletion responses are evaluated using five matched diagnostics: gene-identity increment testing, universal-responsiveness adjustment, tokenization-coverage gating, library-size control, and de-circularized cell-state analysis. Readouts are considered biologically interpretable only if they remain informative after these controls.

**Table 1.**
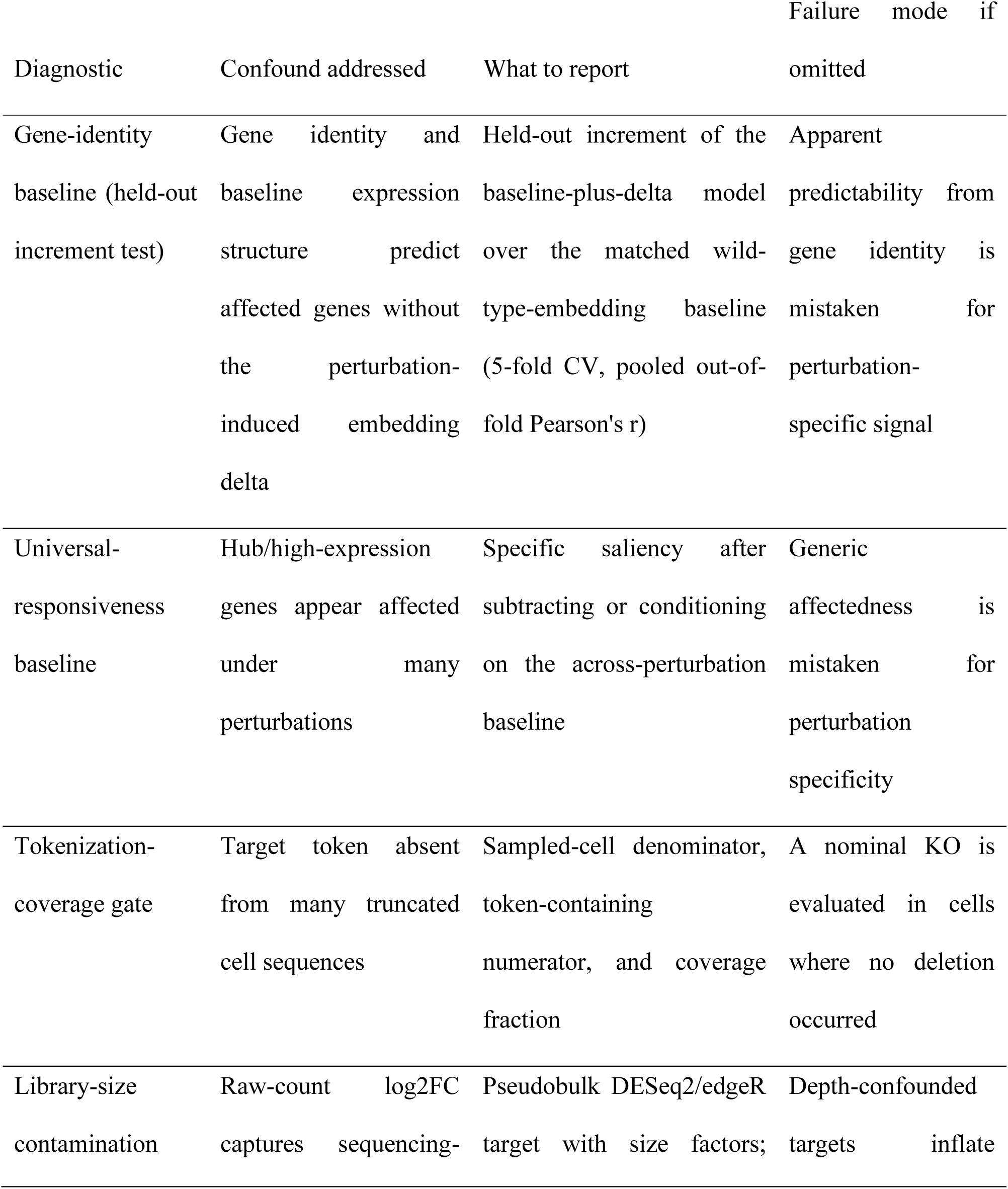
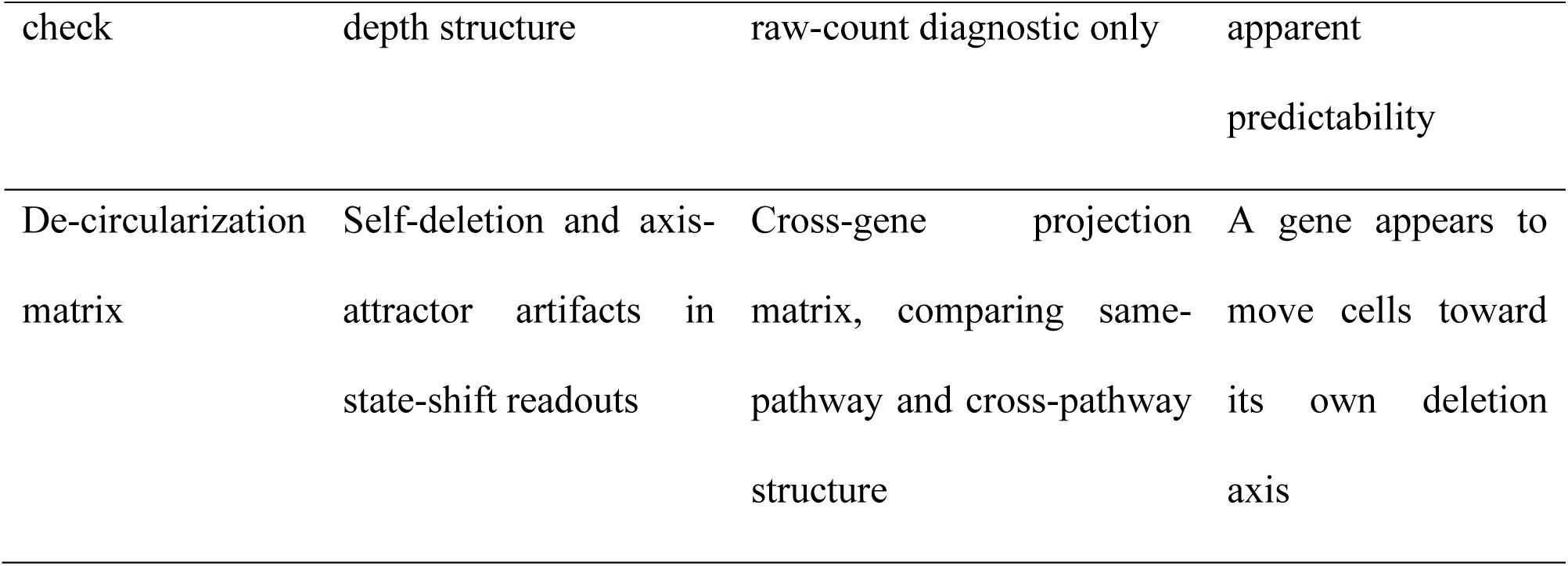
Confound-diagnostic controls and reporting criteria for native in silico perturbation readouts.

We first asked whether the native embedding delta improved prediction of experimentally measured perturbation effects beyond gene identity. In the Frangieh Perturb-CITE-seq dataset, we evaluated CCND1 knockout using a DESeq2-derived pseudobulk differential-expression target. The unperturbed Geneformer embedding of each evaluated gene served as the baseline, and the matched baseline-plus-delta model additionally included the 768-dimensional knockout embedding delta.

The gene-identity baseline achieved a held-out Pearson correlation of r=0.487. Adding the embedding delta yielded an increment of Δr=−0.004. Similar near-zero increments were observed for ridge regression, a regularization-swept multilayer perceptron, and gradient-boosted trees (Δr=−0.004, −0.024, and +0.010, respectively; **Figure 2A**). Thus, across linear and nonlinear readout models, the native embedding delta did not provide a reproducible held-out improvement beyond the matched gene-identity baseline.

**Figure 2.**
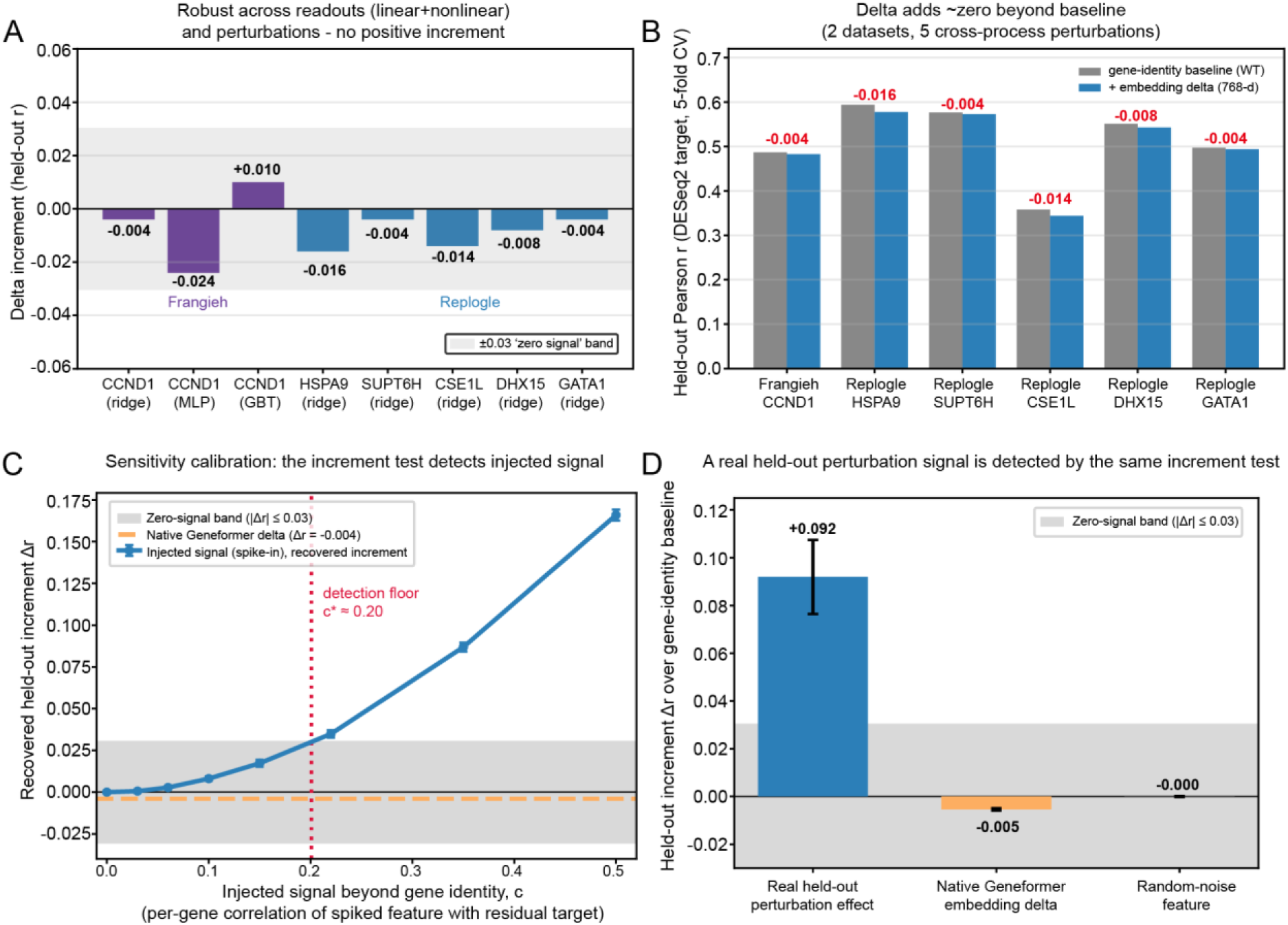
Native embedding-delta increments remain near zero, below the sensitivity of an increment test validated by positive controls. (A) Held-out increments after adding the embedding delta to the gene-identity baseline. The shaded band (∣Δr∣≤0.03) is a visualization threshold, not an equivalence bound. (B) Gene-identity baseline and baseline-plus-delta performance across Frangieh CCND1 and five Replogle perturbations, showing no reproducible positive increment. (C) Signal-injection calibration. Detectable increments emerged at c^∗^≈0.20, whereas the native delta (Δr=−0.004) remained within the near-zero band, indicating that any additional signal was below the detection threshold. (D) Real-signal positive control. A real, held-out CCND1 effect (log2 fold change from a disjoint half of the CCND1 cells) added through the identical increment machinery produced a clear positive increment (Δr = +0.092 ± 0.016 over 20 splits), whereas the native embedding delta (Δr = −0.005) and a random-noise feature (Δr = −0.000) remained within the zero-signal band. The test thus detects real perturbation signal, and the native delta behaves like the noise control.

### Embedding-delta increments remain near zero across datasets and perturbation programs

We next examined whether this result generalized to a more heterogeneous perturbation dataset. Among perturbations represented by at least 50 cells, the mean pairwise perturbation-profile correlation was +0.06 in the Replogle K562 CRISPRi dataset, compared with +0.40 in the Frangieh dataset, indicating greater diversity among Replogle transcriptional responses. Split-half reliability for the selected Replogle perturbations ranged from 0.88 to 0.99, confirming that these perturbation programs were reproducible.

Despite this heterogeneity, embedding-delta increments on DESeq2-derived targets remained near zero across perturbations representing distinct biological processes (**Figure 2B**). The increments were −0.016 for HSPA9, −0.004 for SUPT6H, −0.014 for CSE1L, and −0.008 for DHX15. Each perturbation was supported by 35–46 pseudobulk replicates, compared with 48 controls, and showed strong agreement between DESeq2 and size-factor-normalized fold-change directions.

GATA1 was included as an additional, biologically informative case. Although GATA1 knockdown produced the strongest transcriptional program among the evaluated perturbations, with 766 differentially expressed genes, its embedding-delta increment was also near zero (Δr=−0.004). Because only three pseudobulk replicates were available and the sign-check value was +0.66, GATA1 was considered supportive rather than primary evidence.

These findings argue against dataset homogeneity or weak experimental perturbation effects as explanations for the near-zero increments. The Replogle dataset contained diverse and reproducible perturbation-specific transcriptional programs, yet the native Geneformer embedding delta did not recover detectable information beyond gene identity.

Finally, we calibrated the sensitivity of the held-out increment test by introducing features with known residual signal beyond the gene-identity baseline. A detectable increment emerged when the injected feature had a per-gene correlation of approximately c^∗^=0.20 with the residual target. By contrast, the native delta increment remained within the prespecified near-zero band (Δr=−0.004; **Figure 2C**). Under this calibration, any perturbation-specific information carried by the native embedding delta was therefore below the detection threshold of the analysis.

As a complementary positive control, we confirmed that the increment test recovers genuine perturbation signal, not only synthetic injections. A real, held-out measurement of the CCND1 effect—computed on a disjoint half of the CCND1 cells and passed through the identical machinery—produced a clear positive increment (Δr = +0.092 ± 0.016), whereas the native embedding delta and a random-noise feature both remained within the near-zero band (Δr = −0.005 and −0.000, respectively; **Figure 2D**). Thus the native delta behaves like the noise control rather than like a real perturbation feature, indicating that the near-zero native increment reflects an absence of signal rather than an insensitive test.

### Matched controls identify confounds underlying apparently informative readouts

We next examined exploratory readouts that initially appeared informative but were not intended as primary evidence of perturbation-specific signal. These analyses served as diagnostic stress tests to determine whether the observed signals persisted after matched confound controls.

Directional and scalar readouts were sensitive to score construction and target definition. An oracle-direction diagnostic yielded a correlation of r=+0.39, but the scoring direction was defined using the target itself; reversing that direction reversed the sign of the correlation by construction. A simple readout based on response magnitude and projection achieved r=+0.27 against a raw-count log2 fold-change target, but only r=+0.01 against the primary DESeq2-derived target. Thus, the apparent performance was largely dependent on raw-count structure rather than a robust perturbation-specific signal.

A 768-dimensional embedding-delta probe similarly achieved r=+0.69 on the exploratory target. However, the unperturbed gene-embedding baseline performed comparably, and adding the embedding delta produced no detectable held-out improvement over this matched baseline (**Figure 3A**). The apparent predictability was therefore attributable primarily to gene identity and baseline expression structure rather than to information introduced by token deletion.

**Figure 3.**
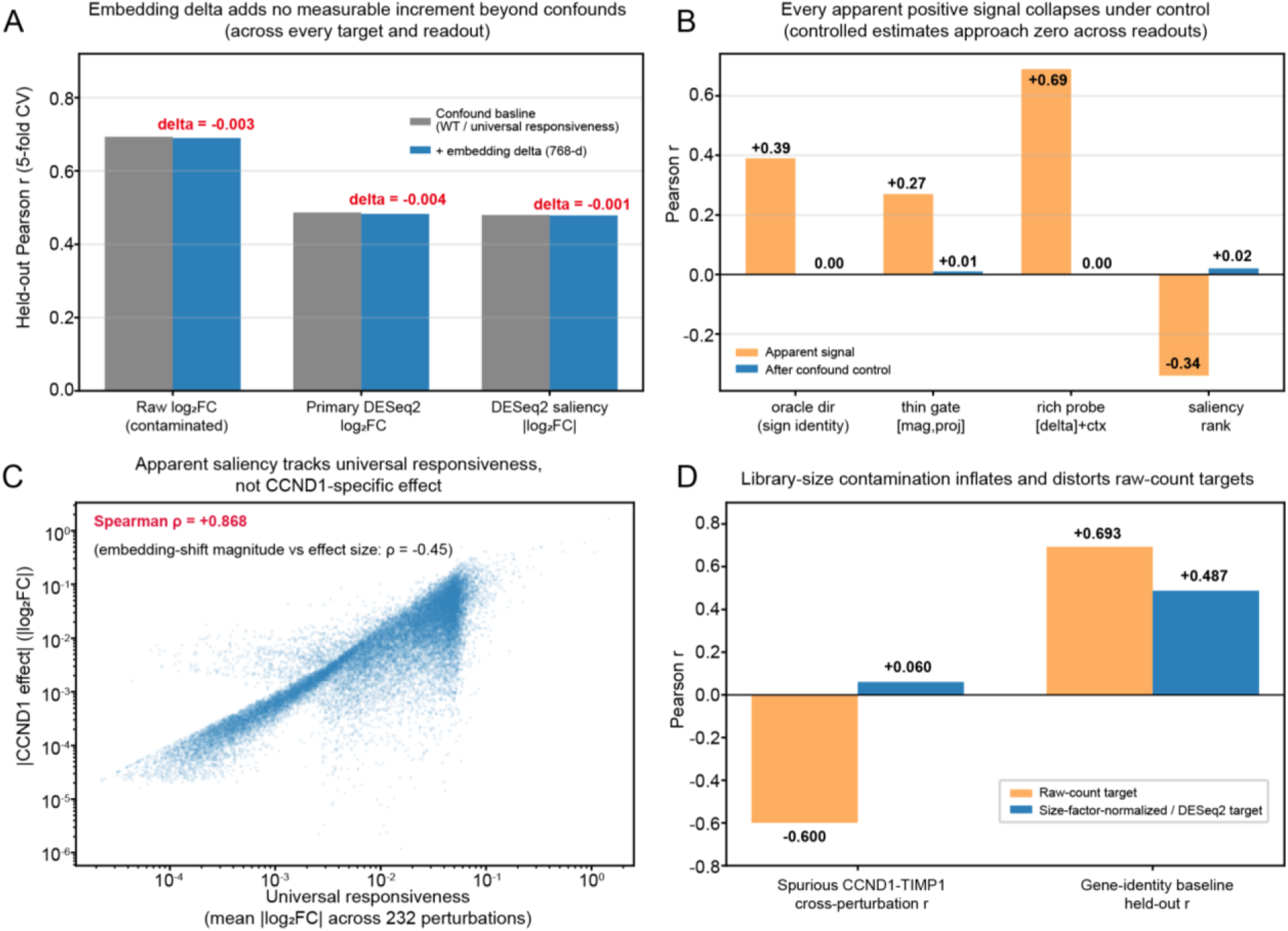
Matched controls attenuate apparent signals and reveal underlying confounds. (A) Adding the 768-dimensional knockout embedding delta provides no held-out improvement over matched baselines for differential-expression and saliency readouts. (B) Apparently informative exploratory readouts are strongly attenuated after controls for score construction, gene identity, library-size structure, and universal responsiveness. (C) Across 232 Frangieh perturbations, universal responsiveness correlates strongly with the absolute CCND1 effect (ρ=0.868), whereas embedding-shift magnitude is inversely associated with biological effect size (ρ=−0.45). (D) Size-factor normalization removes raw-count structure: the CCND1–TIMP1 correlation shifts from r=−0.600 to r=0.060, and gene-identity baseline performance decreases from r=0.693 to r=0.487 across 13,054 genes.

Affected-gene saliency also reflected broad responsiveness. The initial saliency readout correlated with the CCND1 effect at r=−0.34, but the partial correlation after adjustment for universal responsiveness was only r=+0.02. This attenuation indicates that the saliency pattern largely captured genes that respond across many perturbations rather than genes specifically affected by CCND1 knockout.

Together, these matched controls substantially attenuated the apparently informative exploratory readouts. Directional performance depended on target-defined orientation, raw-count performance disappeared after normalized differential-expression analysis, high-dimensional probes did not improve over gene identity, and saliency was removed by adjustment for universal responsiveness (**Figure 3B**).

### Universal responsiveness and library-size structure drive exploratory signal inflation

Across 232 Frangieh perturbations, each gene’s universal mean absolute response was strongly associated with the absolute CCND1 effect (Spearman’s ρ=+0.868). In contrast, embedding-shift magnitude was inversely associated with biological effect magnitude (ρ=−0.45; **Figure 3C**). These findings indicate that apparent affected-gene saliency was dominated by broadly responsive genes and that larger embedding shifts did not preferentially correspond to larger biological effects.

Raw-count fold-change targets also contained substantial library-size structure. The CCND1– TIMP1 correlation shifted from r=−0.600 for raw-count log2 fold change to r=+0.060 after size-factor normalization, while the control-split association approached zero. Library-size structure also inflated apparent predictability: across 13,054 genes, the gene-identity baseline achieved a held-out correlation of r=0.693 on the raw-count target, compared with r=0.487 on the DESeq2-derived target (**Figure 3D**).

These results show that raw-count fold-change targets can inflate apparent model performance and are unsuitable as primary reference targets in this setting. Perturbation effects should instead be estimated using size-factor-normalized pseudobulk differential-expression analyses with appropriate control groups.

### De-circularization does not support pathway-specific cell-state movement

We also evaluated whether native Geneformer knockout shifted cell embeddings toward experimentally observed perturbation states. In the Replogle K562 dataset, in silico GATA1 knockout initially appeared to move cells toward the experimental GATA1-knockdown state, with z≈+4.6 relative to a small random-gene null (n=10).

To reduce self-referential scoring, we constructed a cross-gene projection matrix in which the in silico knockout of gene A was projected onto the experimental perturbation-state axis of a different gene B. This design separated the deleted gene from the axis used for evaluation. Deterministic comparisons among experimentally perturbed genes served as the primary analysis, while the seed-fixed random-gene null was used as a sensitivity check.

After de-circularization, the projection matrix showed no evidence of the expected pathway-specific structure. If native knockout induced pathway-directed state movement, same-pathway off-diagonal projections would be expected to exceed cross-pathway projections. Instead, cross-pathway projections were larger than same-pathway projections (16.7×10^−3^ versus 3.7×10^−3^), opposite to the expected pattern. The same-pathway signal was also small and only weakly separated from the random-gene null (n=6). These findings suggest that the initial GATA1 state shift was more consistent with self-referential scoring than with pathway-specific state movement. Gene groups, projection axes, and null genes are listed in **Supplementary Table S1**.

This cell-level analysis is distinct from the per-gene differential-expression increment test, in which GATA1 likewise showed no detectable embedding-delta increment.

### Tokenization coverage constrains perturbation applicability and estimate stability

We next examined tokenization coverage, defined as the fraction of cells in which the target-gene token was present in the top-4096 input sequence. When the target token was absent, no deletion occurred and the cell did not contribute to the native knockout estimate. Across 217 in-vocabulary Frangieh targets, median coverage was 47%. Several biologically relevant targets had substantially lower coverage, including JAK2 at 2.3% and CD274 at 11% (**Figure 4A**), indicating that native token deletion was applicable to only a small fraction of cells for some perturbations.

**Figure 4.**
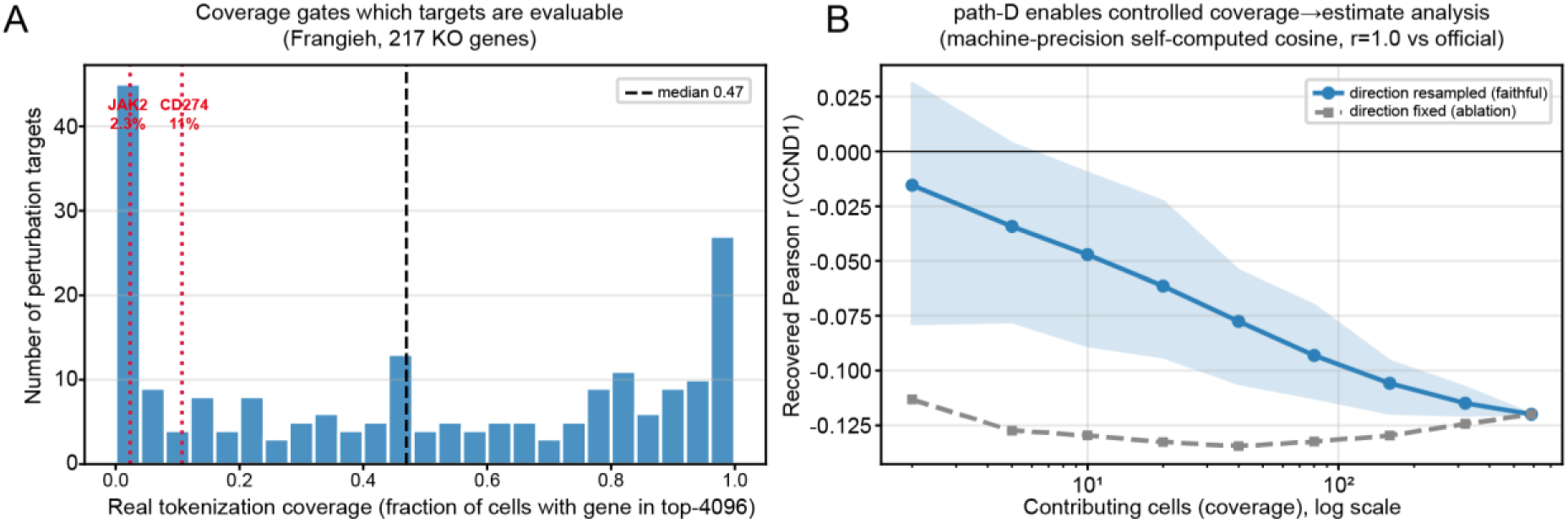
Tokenization coverage constrains perturbation applicability and estimate stability. (A) Distribution of tokenization coverage across 217 in-vocabulary Frangieh targets, defined as the fraction of sampled cells containing the target token in the top-4096 input sequence. JAK2 (2.3%) and CD274 (11%) illustrate low-coverage targets for which native deletion applies to only a small fraction of cells. (B) Controlled subsampling with the numerically matched path-D implementation shows that the recovered CCND1 readout changes with the number of contributing cells and converges as this number increases. Coverage therefore affects applicability and stability but does not establish biological specificity.

The numerically matched path-D implementation enabled direct evaluation of coverage effects. We repeatedly subsampled token-containing cells while holding the model checkpoint, tokenization, target gene, and perturbation operator fixed, thereby isolating the effect of the number of contributing cells.

For the exploratory raw-count CCND1 target, the recovered knockout estimate changed systematically with the number of contributing cells and converged as this number increased (**Figure 4B**). However, convergence produced a stable estimate of the same exploratory readout and did not establish perturbation specificity. An ablation comparing fixed-direction and fully resampled estimates further indicated that the coverage dependence arose primarily from the directional component of the native response.

These results identify tokenization coverage as both an applicability and an estimate-stability constraint. Higher coverage allows native deletion to be applied in more cells and improves estimate stability, but it does not establish biological specificity. Even high-coverage readouts therefore require controls for gene identity, universal responsiveness, library-size structure, and circular state scoring.

## Discussion

This study presents a reusable confound-diagnostic workflow for evaluating native foundation-model in silico perturbation readouts. Using frozen Geneformer token deletion as a case study, we found that the native embedding delta provided no reproducible held-out improvement beyond a gene-identity baseline across two perturbation datasets and linear and nonlinear readout models. Signal-injection calibration confirmed that the increment test could recover added residual signal, whereas the native delta remained below the detection threshold of the analysis. Matched diagnostics further showed that apparently informative readouts could arise from gene identity, universal responsiveness, library-size structure, and self-referential state scoring. Tokenization coverage imposed an additional constraint on perturbation applicability and estimate stability but did not establish biological specificity.

The primary evidence came from held-out increment testing. Adding the 768-dimensional knockout embedding delta to unperturbed gene embeddings did not improve prediction of experimentally measured perturbation effects. This result was consistent across ridge regression, multilayer perceptrons, and gradient-boosted trees and across the Frangieh Perturb-CITE-seq and Replogle K562 CRISPRi datasets. Importantly, the Replogle dataset contained heterogeneous and reproducible perturbation-specific transcriptional programs, arguing against weak or homogeneous benchmark signal as the explanation for the near-zero increments. Moreover, the signal-injection analysis recovered detectable increments once the injected feature carried sufficient residual information, whereas the native delta remained within the near-zero band. Thus, under the tested design, any perturbation-specific information carried by the native embedding delta was below the detection threshold rather than obscured by an insensitive evaluation.

The exploratory analyses identified several distinct mechanisms of apparent signal inflation. Unperturbed gene embeddings supported substantial predictability without information from the knockout operation, indicating that gene identity and baseline expression structure can be mistaken for perturbation-specific signal. Affected-gene saliency largely reflected genes that responded broadly across many perturbations rather than specifically to CCND1 knockout. Raw-count fold-change targets contained library-size structure that inflated apparent predictability and cross-perturbation correlations. Cell-state-shift readouts became misleading when the deleted gene also contributed to the experimental axis used for scoring; after cross-gene de-circularization, the expected pathway-specific structure was not observed. These findings emphasize that apparently strong correlations are insufficient unless the corresponding alternative explanations are explicitly controlled.

Tokenization coverage represents a related but conceptually distinct limitation. Native deletion can occur only in cells whose truncated input sequence contains the target token. Across Frangieh targets, coverage varied widely, and several biologically relevant genes were represented in only a small fraction of cells. Controlled subsampling showed that the recovered estimate changed systematically with the number of contributing cells and converged as that number increased. However, convergence stabilized the evaluated readout without demonstrating that it was biologically specific. Coverage should therefore be treated as an applicability and stability criterion rather than as evidence supporting biological validity.

The contribution of this study is not an additional aggregate benchmark of perturbation prediction, but a framework for determining why a native perturbation readout appears informative and whether that signal persists after matched controls. The numerically matched path-D implementation enables controlled manipulation of the number of contributing cells while holding the model, tokenization, target gene, and perturbation operator fixed. Held-out increment testing directly assesses whether the embedding delta adds information beyond gene identity, whereas universal-responsiveness adjustment, library-size diagnostics, and de-circularized projections address complementary sources of signal inflation. Although tokenization coverage is specific to token-based models, other architectures will require analogous applicability checks to verify that the perturbation operator meaningfully alters the relevant model input or representation.

The empirical scope of the study remains intentionally limited. We evaluated frozen Geneformer native token deletion and did not test fine-tuned perturbation models, decoder-based architectures, inference-time steering, or alternative perturbation operators. The findings therefore should not be generalized to all single-cell foundation models or perturbation-modeling strategies. The cell-state analysis used K562 perturbation axes rather than an independent differentiation or functional-state reference, and the GATA1 analysis had limited pseudobulk replication. In addition, the coverage-to-estimate analysis assessed estimate stability using an exploratory readout and did not establish biological accuracy. These limitations define the scope of the conclusions but do not change the central result: under the tested conditions, the native Geneformer embedding delta provided no detectable perturbation-specific information beyond gene identity.

Native token deletion remains computationally simple and potentially useful as a hypothesis-generating operation, but deletion of an input token should not be assumed to reproduce biological loss of function. Future methods may improve perturbation modeling through supervised fine-tuning, decoder-based response prediction, inference-time steering^14^, rank-aware evaluation^15^, or perturbation operators designed to better approximate biological intervention. Regardless of architecture, foundation-model perturbation readouts should be interpreted only after controlling for gene identity, broad responsiveness, library-size effects, perturbation applicability, and circular evaluation. More broadly, the framework presented here provides practical safeguards for developing, benchmarking, and interpreting foundation-model in silico perturbation methods.

## Author contributions

R.Q. conceived and designed the study, developed the methodology and software, performed the analyses, interpreted the results, and wrote the manuscript. M.M.Z. contributed to data analysis, code review, result interpretation, and manuscript editing. All authors reviewed and approved the final manuscript.

## Declaration of interests

The authors declare no competing interests.

## Supplemental information

Supplemental information includes Supplementary Table S1, which lists the de-circularization gene groups, projection axes, and random-null genes, and Supplementary Table S2, which provides the reported-numbers provenance.

## STAR Methods

### Resource availability

#### Lead contact

Further information and requests for resources should be directed to the lead contact, Ru Qiu.

#### Materials availability

This study did not generate new biological materials.

#### Data and code availability

All datasets analyzed in this study are publicly available as preprocessed scPerturb h5ad files^16^. The analysis code is publicly available as isp-confound-toolkit (https://github.com/willow0077/isp-confound-toolkit) and archived on Zenodo (https://doi.org/10.5281/zenodo.20743523), organized under scripts/ — pipeline/ (path-D caching, resampling, state-shift, de-circularization matrix) and diagnostics/ (token coverage, confound checks, DESeq2 ground truth, increment tests, heterogeneity pre-check, figure generation). The repository includes a reported-numbers provenance table, panel-level figure-reproduction notes, a data/local-path manifest, and supplementary audit notes. The cached path-D per-cell response files, DESeq2 outputs, figure inputs, and audit tables enabling pure-NumPy reproduction of the main figures and increment tests are available on Zenodo: https://doi.org/10.5281/zenodo.20729460.

## Method details

### Datasets and preprocessing

The Frangieh dataset^17^ contains patient-derived melanoma cells and autologous tumor-infiltrating lymphocytes profiled by Perturb-CITE-seq after CRISPR-Cas9 knockout. The scPerturb-harmonized h5ad file used in this study includes cells from control, IFN-γ stimulation, and co-culture conditions, with perturbed genes and a non-targeting control.

The Replogle dataset^18^ contains K562 CRISPR interference perturbations from essential-scale and genome-scale Perturb-seq experiments. The scPerturb-harmonized subset used in this study includes perturbed genes and a non-targeting control.

Raw counts from both datasets were used only as input for size-factor-normalized pseudobulk differential expression or as an explicit diagnostic for library-size contamination. Raw-count log2 fold change was not used as the primary experimental reference target.

Different analyses used purpose-defined subsets according to the relevant inclusion criteria. For the Frangieh CCND1 worked example, analysis subsets are listed in **Table 2**. Replogle analyses used analogous per-target subsets. Perturbation subsets were defined separately for universal-responsiveness analysis, heterogeneity analysis, and tokenization-coverage analysis rather than as a single nested filtering scheme.

**Table 2.**
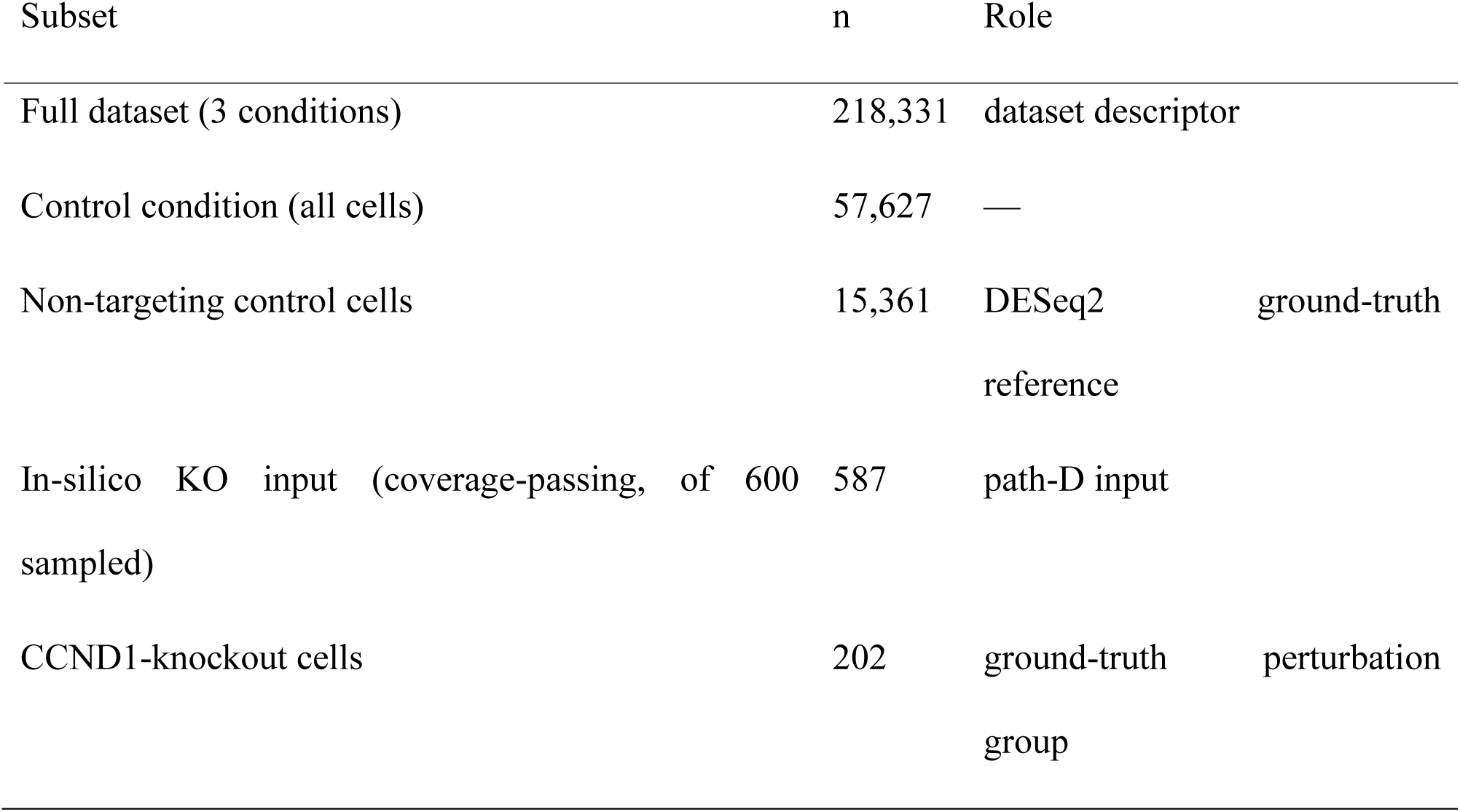
Cell subsets used in the Frangieh CCND1 case study.

### Geneformer model and native in silico knockout

All analyses were performed using the frozen Geneformer V2-104M model without fine-tuning. The model contains 12 transformer layers, a hidden dimension of 768, and a gene-token vocabulary of approximately 20,271 tokens. The specific model checkpoint, Geneformer version, and commit identifier are listed in the Key resources table. Unless otherwise specified, embeddings were extracted from the second-to-last hidden layer (hidden_states[11]; emb_layer = −1 in Geneformer’s convention).

Native in silico knockout was performed by deleting the target-gene token from each cell’s rank-ordered top-4096 token sequence. Wild-type and knockout forward passes were then compared to obtain gene-level or cell-level embedding responses. Gene embeddings were extracted for evaluated genes, whereas cell embeddings were represented by the classification token. Cells in which the target-gene token was absent were not assigned a zero response, because no token deletion occurred in those cells. These cells were excluded from the native knockout estimate.

### Tokenization coverage

Tokenization coverage was defined as the fraction of sampled input cells in which the target-gene token was present in the top-4096 token sequence. The denominator was the total number of sampled cells, and the numerator was the number of cells containing the target token. This definition separates biological cell sampling from perturbation applicability in the tokenized model input. Coverage values were recorded for each target and used to determine whether native token deletion could be meaningfully applied in the sampled cell population.

### Path-D implementation and validation

We implemented the per-cell gene cosine-similarity response to target-gene deletion directly from the frozen Geneformer model and refer to this response as path-D. Validation was performed by comparing path-D outputs with the official Geneformer InSilicoPerturber outputs on the same input cells and target genes. Pearson correlation and maximum absolute error were used as numerical validation metrics. The validation results are reported in the Results.

After validation, per-cell path-D responses were cached and used for downstream NumPy-based resampling analyses. This implementation allowed repeated subsampling of contributing cells without rerunning the transformer model. In the coverage-to-estimate analysis, the number of contributing cells was varied while the model, per-cell tokenization, target gene, and perturbation operator were held fixed. This design isolated the effect of the contributing-cell count from differences among targets, model runs, or perturbation operators.

### Experimental reference differential expression

Primary log2 fold-change targets were computed by pseudobulk aggregation followed by DESeq2^19^ with median-of-ratios size factors, implemented in PyDESeq2^20^; edgeR^21^ provides an equivalent count-based alternative. The exact PyDESeq2 version, DESeq2-compatible settings, and preprocessing scripts are listed in the Key resources table and code repository.

Pseudobulk replicates were defined separately for each dataset to avoid cell-level pseudoreplication^22^. In the Frangieh dataset, pseudobulk replicates were defined by single-guide RNA. In the Replogle dataset, pseudobulk replicates were defined by gemgroup batch. The non-targeting or control population served as the reference group. Log fold changes were estimated with apeglm shrinkage^23^. For each perturbation target, a sign check was performed to confirm that the DESeq2 log fold change agreed in direction with the corresponding size-factor-normalized log2 fold change.

Raw-count log2 fold change was not used as a primary evaluation target. It was used only as an exploratory diagnostic to assess library-size contamination.

### Held-out increment testing

For each readout, we evaluated whether the native knockout embedding delta added predictive information beyond a matched baseline feature set. The primary baseline was the wild-type gene embedding, defined as the unperturbed embedding of each evaluated gene. This baseline captures gene identity and baseline expression structure without using the native in silico knockout response.

For each model class, the baseline feature set was compared with the same feature set plus the 768-dimensional embedding delta. Both feature sets were evaluated using the same model class and the same cross-validation splits. The evaluated readout models included ridge regression, a regularization-swept multilayer perceptron, and gradient-boosted trees. All readout models used a fixed random seed (0) and 5-fold cross-validation (KFold, shuffle enabled, random_state = 0), with feature standardization fit on the training fold only to avoid leakage. Ridge regression used RidgeCV over 13 logarithmically spaced L2 penalties (10⁻² to 10⁴). The multilayer perceptron used a single hidden layer of 32 units with early stopping (max_iter = 400, n_iter_no_change = 10), and its L2 penalty (α) was swept over {1, 10, 30, 100}. Gradient-boosted trees used a histogram-based gradient-boosting regressor (max_depth = 3, L2 regularization = 1.0, early stopping). Full configurations and seeds are provided in the deposited analysis code (scripts/diagnostics/nonlinear_seal.py).

All reported correlations were held-out estimates from 5-fold cross-validation. Out-of-fold predictions were pooled across folds before Pearson’s r was computed. In-sample correlations were not used as evidence of predictive performance. Delta increment was defined as the difference between the held-out performance of the baseline-plus-delta model and that of the corresponding matched baseline model.

Signal-injection calibration. To calibrate the sensitivity of the increment test, a synthetic feature was constructed with a controlled per-gene correlation c to the residual of the DESeq2 target after the gene-identity baseline, and its recovered increment was evaluated under the identical cross-validation. The detection floor was the smallest c whose mean increment exceeded the |Δr| ≤ 0.03 band.

Real-signal positive control. To verify that the increment test can recover genuine perturbation signal, the CCND1 cells were split into two disjoint halves (seeded). The per-gene log2 fold change measured on the first half was used as an added feature, and the log2 fold change measured on the second half served as the held-out target, ensuring non-circularity. This feature was evaluated against the 768-dimensional gene-identity baseline under the identical 5-fold cross-validation and Ridge configuration used for the native delta, alongside the native delta and a random-noise feature as references. Increments were averaged over 20 random splits.

### Universal-responsiveness analysis

Universal responsiveness was used to assess whether affected-gene saliency reflected perturbation-specific response or broad responsiveness across perturbations. For each evaluated gene, a universal-responsiveness score was computed from its mean absolute response across perturbations satisfying the relevant inclusion criteria. Saliency analyses then evaluated whether apparent perturbation-specific signal remained after conditioning on or subtracting this universal-response component.

### Library-size contamination analysis

Library-size contamination was evaluated by comparing raw-count log2 fold-change targets with size-factor-normalized pseudobulk differential-expression targets. Raw-count targets were used only as diagnostics to test whether sequencing-depth structure could inflate apparent predictability or create spurious cross-perturbation correlations. Primary perturbation-effect analyses used size-factor-normalized DESeq2 targets rather than raw-count log2 fold changes.

### De-circularized cell-state-shift analysis

Cell-state-shift analyses evaluated whether an in silico knockout moved cells toward an experimentally observed perturbation state in embedding space. To reduce self-referential scoring, we constructed a de-circularization matrix in which the knockout of gene A was projected onto the perturbation-state axis of a different gene B. This cross-gene projection design separates the deleted gene from the state axis used for scoring.

Same-pathway off-diagonal projections were compared with cross-pathway projections to evaluate pathway-specific state movement after separating the deleted gene from the scoring axis. A seed-fixed random-gene null was used as a sensitivity check. Gene groups, projection axes, and random-null genes are provided in **Supplementary Table S1**.

### Coverage-to-estimate analysis

Coverage-to-estimate analysis was performed using cached path-D responses. For a fixed target gene and perturbation operator, contributing cells were repeatedly subsampled from the token-containing cell set. The recovered native-knockout estimate was recomputed after each subsampling step. This procedure manipulated the number of contributing cells while holding the model, per-cell tokenization, target gene, and perturbation operator fixed. Fixed-direction and faithfully resampled estimates were compared to identify which component of the estimate was most sensitive to coverage.

### Diagnostic workflow

The confound-diagnostic workflow consisted of five matched controls applied across the analyses. The gene-identity baseline tested whether the embedding delta provided held-out incremental information beyond unperturbed gene features. The universal-responsiveness control assessed whether affected-gene saliency reflected perturbation-specific effects rather than broad responsiveness across perturbations. The tokenization-coverage gate determined whether the target-gene token was present in enough sampled cells for native token deletion to be applicable. The library-size contamination check compared raw-count and size-factor-normalized differential-expression targets to identify sequencing-depth artifacts. The de-circularization matrix evaluated whether cell-state-shift readouts remained specific after separating the deleted gene from the perturbation-state axis used for scoring. **Table 1** lists each diagnostic, the corresponding confound, and the quantity reported for each analysis.

### Quantification and statistical analysis

Pearson’s correlation was used to quantify agreement between model readouts and continuous differential-expression targets. Spearman’s correlation was used for rank-based analyses, including universal-responsiveness and perturbation-profile comparisons. For held-out increment testing, Pearson’s r was computed from pooled out-of-fold predictions generated by 5-fold cross-validation. The same cross-validation splits were used for each matched baseline and baseline-plus-delta comparison.

Delta increment was defined as the held-out Pearson’s r of the baseline-plus-delta model minus the held-out Pearson’s r of the matched baseline model. A positive increment indicated additional predictive information from the embedding delta beyond the baseline feature set. Small increments within the prespecified near-zero visualization band were interpreted as no detectable incremental signal rather than as formal evidence of equivalence.

For differential-expression targets, pseudobulk replicates were defined by single-guide RNA in the Frangieh dataset and by gemgroup batch in the Replogle dataset. DESeq2 sign checks were used to verify agreement between shrinkage-based log fold changes and size-factor-normalized log2 fold changes.

For cell-state-shift analyses, deterministic cross-gene projection comparisons were used as the primary de-circularization assessment. Seed-fixed random-gene nulls were used as sensitivity checks. Software versions and package identifiers are listed in the Key resources table; random seeds and model hyperparameters are specified in the deposited analysis code (scripts/diagnostics/nonlinear_seal.py).

## Notes

https://doi.org/10.5281/zenodo.20743523

https://doi.org/10.5281/zenodo.20729460

https://github.com/willow0077/isp-confound-toolkit

### Competing Interest Statement

The authors have declared no competing interest.

