## Supplementary table S1 for "A confound-diagnostic toolkit for in silico perturbation with single-cell foundation models"

**Supplementary Table S1. De-circularization cross-gene projection matrix**

**Parameters**: N_CTRL = 100, N_KO = 100, N_RANDOM = 6, SEED = 0; CLS-token embeddings. Projection values are shown as ×10⁻³. The machine-readable per-cell table (72 cells) is available in the code repository (Supplementary_Table_S1_decircularization_cells.csv).

**Readout**. Each projection axis B is the real-perturbation state axis of a real gene, defined as normalize(mean CLS of that gene's real-KO cells − mean CLS of control cells). The in-silico knockout of gene A is scored as the mean over control cells (that contain the A token) of the projection of the per-cell CLS shift (CLS_KO − CLS_WT) onto axis B. The matrix is therefore directional: M[A][B] ≠ M[B][A].

Diagonal entries M[A][A] carry the self-deletion / circular component and are reported below for transparency but excluded from the specificity comparison; pathway specificity would require same-pathway off-diagonal > cross-pathway off-diagonal (see Methods, “De-circularized cell-state-shift analysis”).

**1. Gene groups**

| Gene | Group | Pathway / role |
| --- | --- | --- |
| HSPA9 | mitochondrial (MITO) | mitochondrial HSP70 chaperone |
| PHB | mitochondrial (MITO) | mitochondrial prohibitin |
| PHB2 | mitochondrial (MITO) | mitochondrial prohibitin |
| GATA1 | non-mitochondrial real | erythroid / lineage transcription factor |
| CSE1L | non-mitochondrial real | nuclear transport |
| SUPT6H | non-mitochondrial real | transcription elongation |

MITO = {HSPA9, PHB, PHB2}. The mitochondrial cluster is the same-pathway group; GATA1, CSE1L, and SUPT6H are three distinct non-mitochondrial pathways used as cross-pathway controls.

**2. Cross-gene projection matrix (×10⁻³)**

Rows are in-silico knockout genes A; columns are the real-KO state axes B of the same six genes (HSPA9, PHB, PHB2, GATA1, CSE1L, SUPT6H). n is the number of control cells containing the A token that contributed to the row.

| KO gene A | Group | n | HSPA9 | PHB | PHB2 | GATA1 | CSE1L | SUPT6H |
| --- | --- | --- | --- | --- | --- | --- | --- | --- |
| HSPA9 | mitochondrial | 97 | −16.0 | −33.0 | −21.0 | 13.8 | −7.7 | −3.3 |
| PHB | mitochondrial | 100 | 16.4 | 9.1 | 9.2 | 61.9 | −28.3 | 38.6 |
| PHB2 | mitochondrial | 92 | 37.6 | 12.8 | 55.6 | 25.2 | 17.3 | 32.4 |
| GATA1 | non-mito real | 68 | −37.7 | −41.6 | −24.4 | 37.9 | −70.7 | 1.2 |
| CSE1L | non-mito real | 81 | 4.8 | −6.9 | 3.4 | 13.4 | −10.9 | 3.0 |
| SUPT6H | non-mito real | 67 | −5.9 | −20.1 | −7.9 | 11.7 | −15.4 | −1.4 |
| NADK2 | random null | 29 | −2.8 | −15.0 | −5.2 | 10.7 | −15.7 | −0.2 |
| CCDC169 | random null | 9 | −0.3 | −5.2 | −3.8 | 14.2 | 3.0 | 7.4 |
| GBF1 | random null | 23 | 1.3 | −6.0 | −0.9 | 3.5 | −9.3 | −1.2 |
| HNRNPH1 | random null | 95 | −4.6 | −15.7 | −7.7 | −1.1 | −3.3 | −6.4 |
| PARS2 | random null | 13 | −12.0 | −31.7 | −18.4 | 10.5 | −19.5 | −6.3 |
| ZNF358 | random null | 9 | −10.7 | −29.4 | −12.3 | 7.3 | −36.7 | −10.1 |

The full per-cell table — knockout_gene_A, projection_axis_B, A_group, B_group, relation, projection_value_x1e-3, n_contributing_cells, included_in_specificity_comparison — is available in the code repository (Supplementary_Table_S1_decircularization_cells.csv).

**Composition of the specificity comparison**

Each of the 72 cells is assigned a relation label; only off-diagonal same- and cross-pathway cells enter the headline comparison.

| Relation | Cells | Included in specificity comparison | Mean (×10⁻³) |
| --- | --- | --- | --- |
| self / diagonal (self-deletion, circular) | 6 | No — excluded | +12.4 |
| same-pathway off-diagonal (MITO → MITO) | 6 | **Yes** | +3.7 |
| cross-pathway off-diagonal (MITO → non-MITO) | 9 | **Yes** | +16.7 |
| real off-diagonal (non-MITO A → any B) | 15 | No — not part of the MITO-anchored comparison | — |
| null: random → MITO axis | 18 | No — defines the complementary null | −10.0 ± 9.0 |
| null: random → non-MITO axis | 18 | No | — |

**3. Random-null genes**

Seed-fixed draw, N_RANDOM = 6, SEED = 0, drawn (without replacement) from in-vocabulary genes not among the six real genes, in scripts/pipeline/state_shift_matrix.py.

| Random-null gene | N_RANDOM | seed | draw index |
| --- | --- | --- | --- |
| NADK2 | 6 | 0 | 1 |
| CCDC169 | 6 | 0 | 2 |
| GBF1 | 6 | 0 | 3 |
| HNRNPH1 | 6 | 0 | 4 |
| PARS2 | 6 | 0 | 5 |
| ZNF358 | 6 | 0 | 6 |

The complementary random-gene null is the projection of these six knockouts onto the three mitochondrial axes (18 values).

**4. Interpretation**

The decisive comparison is deterministic and uses only real genes: cross-pathway projections (+16.7 ×10⁻³) exceed the same-pathway off-diagonal (+3.7 ×10⁻³) — the reverse of the same > cross pattern that pathway specificity requires. The complementary same-pathway-vs-null test is only weakly separated (z ≈ +1.51) and is treated as supporting, not load-bearing; diagonal self-deletion entries are excluded. The apparent GATA1 cell-state shift is therefore consistent with a self-deletion / axis-attractor artifact rather than pathway-specific state movement.

**Caveat.** This analysis uses a Replogle K562 knockout-state proxy, not a true differentiation / effector axis; it tests de-circularized pathway specificity, not biological state-transition prediction.
